# Thiol-based mucolytics exhibit antiviral activity against SARS-CoV-2 through allosteric disulfide disruption in the spike glycoprotein

**DOI:** 10.1101/2021.07.01.450701

**Authors:** Yunlong Shi, Ari Zeida, Caitlin E. Edwards, Michael L. Mallory, Santiago Sastre, Matías R. Machado, Raymond J. Pickles, Ling Fu, Keke Liu, Jing Yang, Ralph S. Baric, Richard C. Boucher, Rafael Radi, Kate S. Carroll

## Abstract

Small molecule therapeutics targeting severe acute respiratory syndrome coronavirus 2 (SARS-CoV-2) have lagged far behind the development of vaccines in the fight to control the COVID-19 pandemic. Here, we show that thiol-based mucolytic agents, P2119 and P2165, potently inhibit infection by human coronaviruses, including SARS-CoV-2, and decrease the binding of spike glycoprotein to its receptor, angiotensin-converting enzyme 2 (ACE2). Proteomics and reactive cysteine profiling link the antiviral activity of repurposed mucolytic agents to the reduction of key disulfides, specifically, by disruption of the Cys379–Cys432 and Cys391–Cys525 pairs distal to the receptor binding motif (RBM) in the receptor binding domain (RBD) of the spike glycoprotein. Computational analyses provide insight into conformation changes that occur when these disulfides break or form, consistent with an allosteric role, and indicate that P2119/P2165 target a conserved hydrophobic binding pocket in the RBD with the benzyl thiol warhead pointed directly towards Cys432. These collective findings establish the vulnerability of human coronaviruses to repurposed thiol-based mucolytics and lay the groundwork for developing these compounds as a potential treatment, preventative and/or adjuvant against infection.

## Introduction

Control of pandemics requires rapid and sensitive testing, widely available vaccines, and effective therapeutic agents for the infected. For the current SARS-CoV-2 pandemic, impressive progress in the development and testing of vaccines has been achieved. However, the development of effective SARS-CoV-2 antiviral therapeutics has lagged. Multiple strategies have been employed to identify effective anti-SARS-CoV-2 therapeutic agents. Classic antiviral strategies that focused on viral replication and propagation, *e.g*., protease and RNA polymerase inhibitors, have received much attention but have produced limited clinical efficacy to date.^1-5^ Many therapeutic strategies have focused on the intricate SARS-CoV-2 cellular entry processes that include interactions of the spike glycoprotein’s receptor-binding domain (RBD) with the angiotensin-converting enzyme 2 (ACE2), cell surface proteases, and complex fusion events.^6,7^ The development of antiviral agents that target the receptor binding and entry processes requires precise knowledge of the dynamic structures of the viral elements that mediate the binding and entry of virus into target cells.^8,9^

Disulfide bond formation is central to the dynamic structure of many viral receptor binding and entry/fusion proteins.^10^ The role of disulfide bonds in cognate receptor binding proteins, *i.e*., spike proteins, has been widely studied in coronaviruses, including mouse hepatitis virus (MHV), SARS-CoV, and SARS-CoV-2.^11-13^ Viral disulfides are initially formed in the endoplasmic reticulum (ER). These bonds, important for both virus binding and fusion, are further stabilized by the oxidizing extracellular milieu.^10,13^ The SARS-CoV-2 RBD contains a four disulfide pairs: 1) Cys480–Cys488 situated at the ACE2 binding surface, and 2) Cys336–Cys361, Cys379–Cys432, Cys391–Cys525 to stabilize the β sheet structure.^8^ The position of these disulfides in RBD crystal structures, has led to speculation that reduction of these bonds may have untapped therapeutic utility.^14-18^ However, the precise function of these disulfide pairs cannot be read from structure alone, and cysteine reactivity mapping has not been performed to investigate this hypothesis.

Thiol-based mucolytic agents, often administrated orally or as aerosols, are currently in use as therapeutics for pulmonary diseases, *e.g*., cystic fibrosis.^19-21^ Of these agents, preclinical compounds P2119 and P2165 (Figure 1a), originally identified from a library of glycosylated thiols and advanced as mucolytics,^22,23^ are unique as they are restricted to the extracellular space, *i.e*., not membrane permeant, P2119 and P2165 also exhibit a greater therapeutic index compared to clinically available therapeutics, like *N*-acetylcysteine (NAC)^23^ which is taken up by the cell and metabolized to hydrogen sulfide (H_2_S)^24^ a second messenger that initiates pleiotropic changes in myriad targets.^25^ P2119 and P2165 also afford an opportunity to compare the intrinsic potency of a monothiol versus a dithiol compound, and their associated mechanisms of action (Figure 1b).

**Figure 1.**
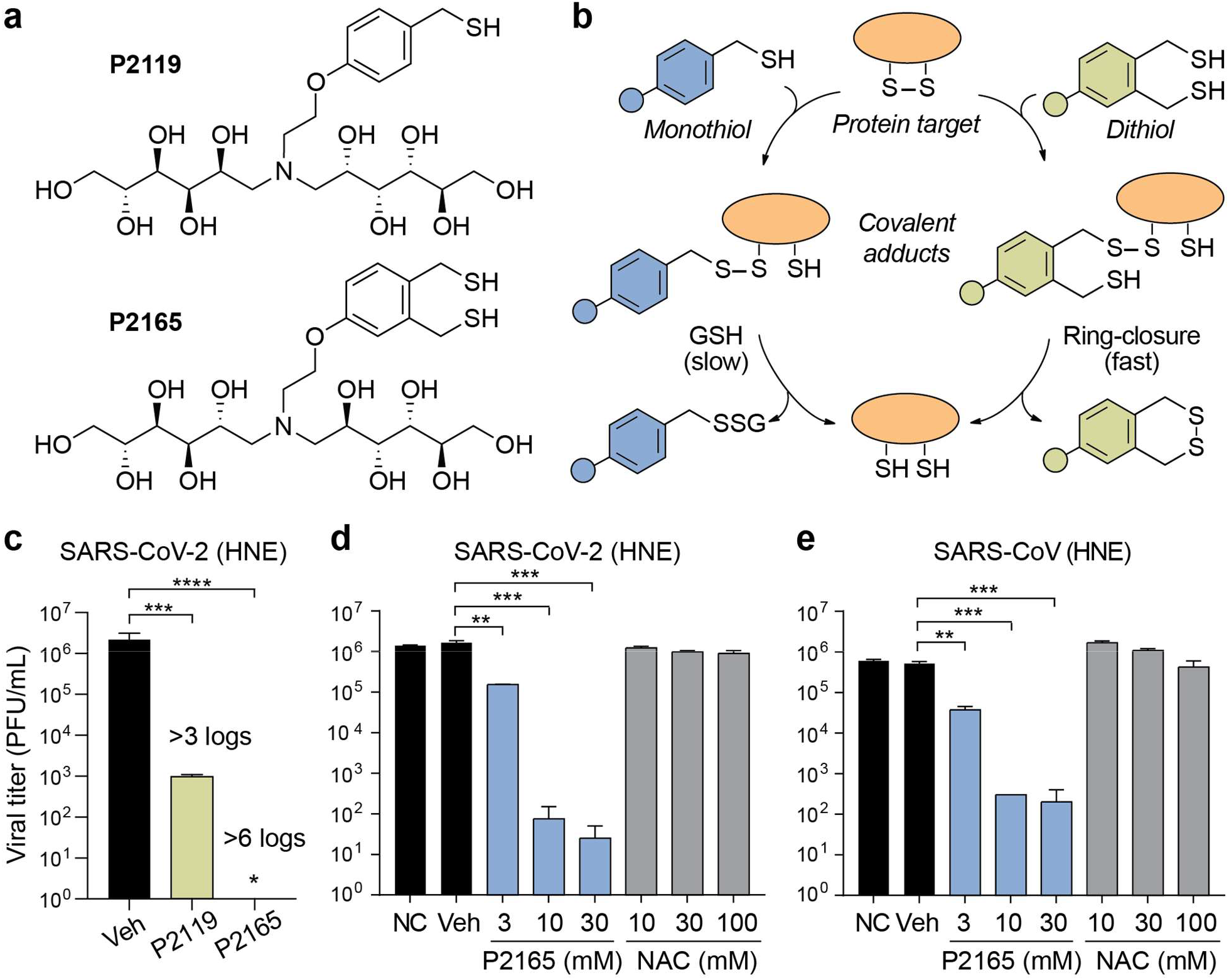
Thiol-based mucolytic compounds have potent virucidal activity against human coronaviruses. **a**, Chemical structures of P2119 and P2165 mucolytic agents. **b**, Mechanisms of disulfide reduction by thiol-based mucolytics. Adducts formed with P2119 require a second thiol equivalent, e.g., from GSH or drug, to resolve the mixed disulfide, whereas the adduct formed with P2165 rapidly undergo intramolecular thiol-disulfide exchange. **c-e**, Comparison of 72-h titers between non-treated control (NC), PBS vehicle (Veh), or mucolytic-treated SARS-CoV-2 D614G infected primary human nasal epithelial (HNE) cultures at an MOI of 0.1. Triplicated titers of the virus in cultures from the same donor were analyzed by paired t tests. ** P < 0.01; ***P < 0.001; ****P < 0.0001.

Here, we show that thiol-based mucolytic agents, P2119 and P2165, potently inhibit infection by human coronaviruses, including SARS-CoV-2, and decrease the binding of spike glycoprotein to its receptor, ACE2. The antiviral activity of repurposed mucolytic agents is linked to the reduction of disulfides in the RBD of spike glycoprotein. Proteomics and reactive cysteine mapping shows that the Cys480–Cys488 pair, located in the loop region of the ACE2 binding surface, is not susceptible to alkylation during live cell infection, establishing the stability of this disulfide in a native setting. By contrast, Cys432 and Cys525, which form disulfides with C379 and Cys391, respectively, are identified as hyper-reactive cysteines that form semi-stable disulfides. Molecular docking analysis provides insight into the targeting of Cys432 and Cys525 by mucolytics. These simulations predict that reduction of these three disulfides control ACE2 binding by triggering conformational changes in the RBD. The latter finding suggests that Cys379–Cys432 and Cys391–Cys525 bonds are allosteric, a unique and rare category of disulfide, distinct from structural and catalytic roles. This work establishes the vulnerability of human coronaviruses to repurposed thiol-based mucolytics, laying the groundwork to develop these compounds as a potential treatment, preventative and/or adjuvant against infection.

## Results

### Thiol-based mucolytic agents have potent antiviral activity against human coronaviruses

The antiviral activities of P2119 and P2165 were first evaluated in a recombinant infectious clone of SARS-CoV-2 virus produced in human nasal epithelial (HNE) cells (Figure 1c).^26^ Exposure of SARS-CoV-2 to monothiol P2119 or dithiol P2165 mucolytics at 30 mM (note: this concentration was chosen based on mucolytic activity and effective surface airway deposition in mouse lungs^22^) resulted in respective >3 log and >6 log reductions in viral titer. To contextualize these findings, we compared dose-dependent virucidal activity of NAC, the gold-standard for approved care in thiol-based molecule mucolytics, and P2165 against SARS-CoV-2 (Figure 1d). Dose-dependent inhibition by P2165 was noted at concentrations over 3 mM (1-3 logs), while NAC was ineffective as a virucidal agent at concentrations up to 100 mM. Analogous virucidal activities were observed for the closely related coronavirus SARS-CoV (Figure 1e), which shares ∼75% identity with SARS-CoV-2 spike protein and three of four conserved disulfides in the RBD.^27^ Coronavirus NL63-CoV^28^ also utilizes ACE2 as its receptor^29^ and has RBD disulfides that are essential for protein stability and/or receptor binding.^30^ Exposure of NL63-CoV to P2119 (10 or 30 mM) or P2165 (10 mM) and infection of LLC-MK2 cells showed similar reduction of viral titer (>3-4 logs; Figure S1a). Finally, since glycosylation of the spike protein likely varies with cell type^31^ the activity of thiol-based mucolytics was also tested with SARS-CoV-2 propagated in Vero E6 cells (Figure S1b). Here, P2119 reduced SARS-CoV-2 titers by ∼1 and 3 logs at 10 and 30 mM, respectively. P2165 reduced virus titers by ∼2 and 4 logs at 10 and 30 mM, respectively. These collective data indicate that P2119 and P2165 have potent antiviral activity irrespective of the cell type producing SARS-CoV-2, that the dithiol is a more effective virucidal agent compared to the monothiol, and hint at a common mechanism of action for these thiol-based mucolytics.

### Thiol-based mucolytics inhibit SARS-CoV-2 spike binding to human ACE2 receptor

P2119 and P2165 are *p*-methoxybenzyl thiols conjugated to glucose (P2119) or mannose (P2165) monomers. The sugar units impart hydrophilicity and block diffusion of mucolytic compounds into the cell where viral enzymes, *e.g*., proteases and mRNA polymerase, hijack host machinery. Since P2119 and P2165 are membrane impermeant and reduction of titer can be observed when the virus is exposed to these compounds prior to cell infection, we hypothesized that P2119 and P2165 act by inhibiting viral entry. To examine this possibility, we assessed *in vitro* binding between recombinant spike receptor binding domain (RBD) from SARS-CoV-2 and immobilized human ACE2. In this workflow, spike RBD was exposed to thiol-based mucolytics, subjected to spin gel filtration for small-molecule removal, and tested for ACE2 receptor binding. Compared to vehicle, P2165 and P2119 treatment shifted the half maximal effective concentration (EC_50_) for ACE2-RBD binding by roughly 6-fold and 4-fold, respectively, while the powerful phosphine-based reducing agent, tris(2-carboxyethyl)phosphine (TCEP) completely blocked the binding of RBD to ACE2 (Figure 2a). These data show that reduction of disulfides in spike RBD decreases binding to ACE2 receptor and, by extension, is likely to hamper the ability of this virus to infect host cells. Interestingly, the trend in receptor-spike binding *i.e*., TCEP >> P2165 > P2119, suggests that antiviral activity is correlated to the intrinsic reducing activity of mucolytic agents.

**Figure 2.**
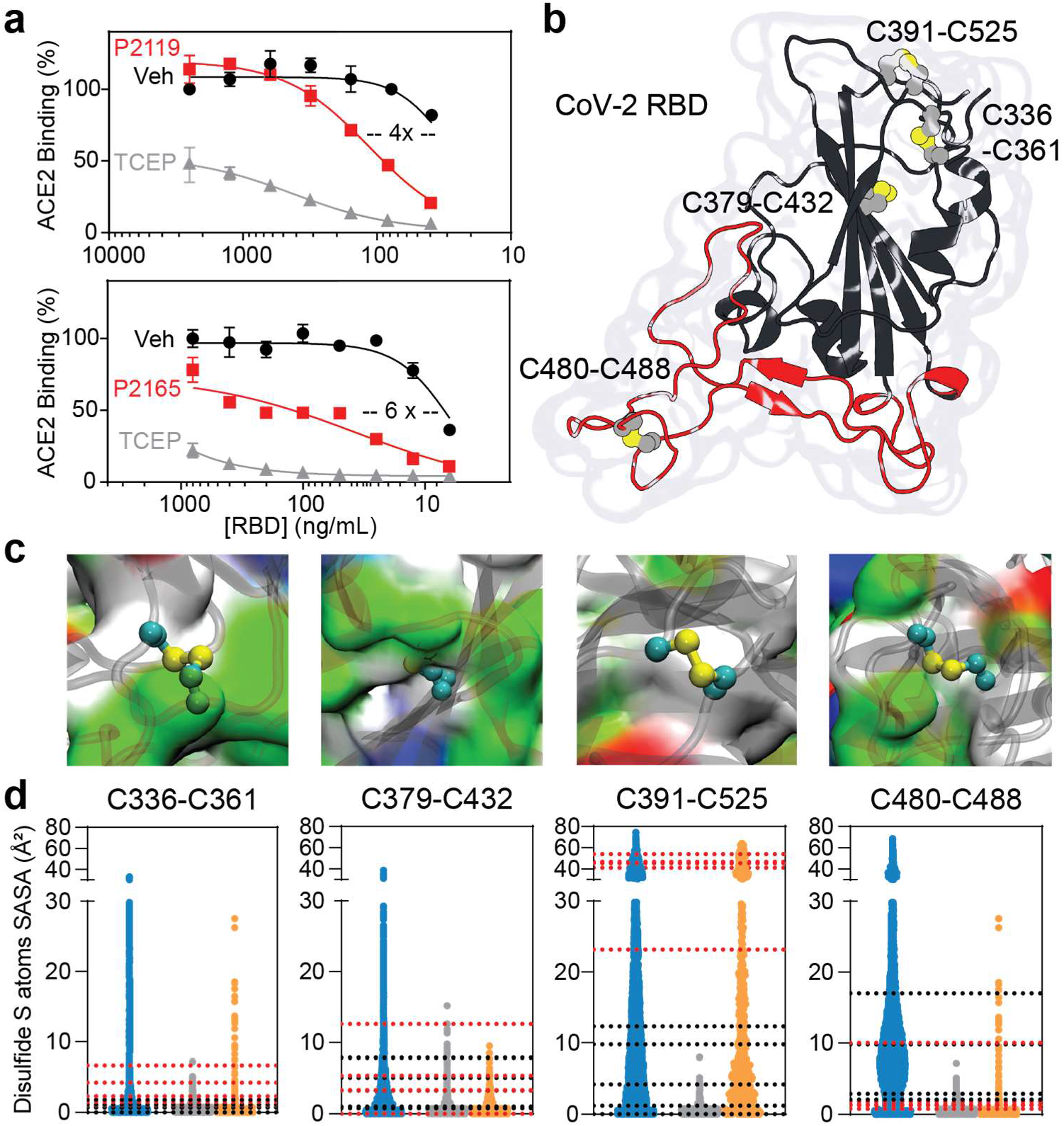
Disulfides of SARS-CoV-2 spike RBD. **a**, Binding of recombinant SARS-CoV-2 spike RBD to immobilized ACE2. Experiments were performed in quadruplicate with vehicle, or with 10 mM P2119, P2165, or TCEP. **b**, Crystal structure of SARS-CoV-2 RBD showing four disulfides and RBM, colored in red. **c**, Protein surface view illustrating the microenvironment of each RBD disulfide and colored by residue type: white, hydrophobic; green, polar; red, acidic; blue, basic. **d**, Solvent accessible surface area (SASA, Å^2^) calculated for RBD disulfides. Blue distributions correspond to isolated RBD MD simulations, gray to averaged three monomers in closed conformation and orange to open conformation from MD simulations performed for the entire spike glycoprotein trimer. Dotted lines correspond to calculated SASA values from different cryo-EM spike glycoprotein structures (black, closed conformations PDB 6XM5, 7KRS, 7BNM; red, open conformations PDB 6XM3, 6XM4, 7KRR, 7BNN, 7BNO).

The globular-shaped SARS-CoV-2 RBD consists of several unstructured or poorly structured regions surrounding a β-sheet domain core and in its native state has four disulfide bonds (Figure 2b and Figure S2). The microenvironment of each disulfide is very different in terms of amino acid sequence and physicochemical properties (Figure 2c). RBD flexibility produces complex dynamic motions for all disulfides, as illustrated by the solvent accessible area (SASA) distributions obtained by molecular dynamics (MD) simulations of either the isolated RBD domain or the entire spike glycoprotein ectodomain in 3-down or 1-up RBD conformations (closed or open) (Figure 2d).^32^ In this analysis, the most buried disulfides were Cys336–Cys361 and Cys379–Cys432, while Cys391–Cys525 and Cys480–Cys488 were the most solvent accessible with isolated and open RBD structures showing comparable SASA values. Notably, similar microenvironment diversity is also captured by comparing spike structures solved under different experimental conditions.

### Mapping mucolytic-sensitive disulfides in human and SARS-CoV-2 cysteinomes in native virus

Based on the promising antiviral studies above, we next sought to identify disulfides that exhibit sensitivity to thiol-based mucolytics in extracellular human and SARS-CoV-2 proteomes. Supernatants from cultured HNE cells infected with SARS-CoV-2 were exposed to P2119 or P2165, subjected to differential alkylation redox proteomics, and quenched by urea (Figure 3a and Table 1, Supporting Information). Cysteine-containing peptides that undergo alkylation by iodoacetamide (IAM) after reduction with P2119 or P2165 represent targets of these thiol-based mucolytics (“mucolytic-sensitive Cys”). In addition, monothiol compounds, like P2119, can form mixed disulfides with cysteines in protein targets (“mucolytic-modified Cys”). whereas dithiols, like P2165, are designed to undergo rapid thiol-disulfide exchange to minimize stable covalent modification by the mucolytic agent.

**Figure 3.**
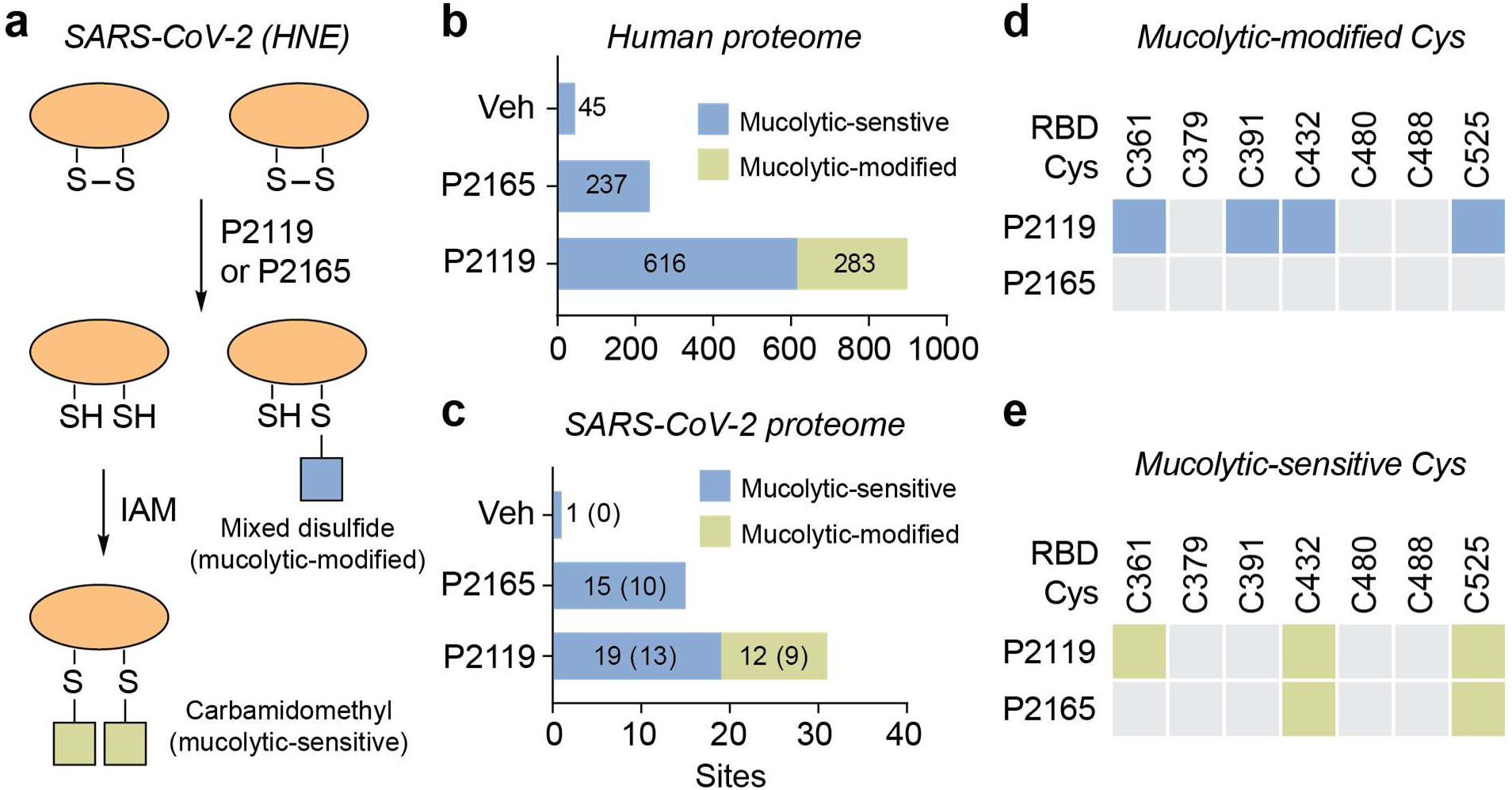
Cysteines in human and SARS-CoV-2 proteomes targeted by mucolytic agents. **a**, Workflow for identifying mucolytic-sensitive and mucolytic-modified cysteines in SARS-CoV-2. HNE cultures were infected with SARS-CoV-2 D614G and diluted in PBS to an MOI of 0.1 on the apical surface. At 96 hpi, apical washes were incubated with PBS vehicle, P2119 (30 mM), or P2165 (30 mM). Samples were desalted, concentrated, alkylated by IAM, and processed using standard proteomic procedures. **b-c**, Number of peptides with mucolytic-sensitive or mucolytic-modified cysteines in human and viral proteomes. Values in parenthesis are number of spike peptides. **d-e**, Mucolytic-modified and mucolytic-sensitive RBD cysteines of native spike. Colored boxes indicate modified or redox-sensitive sites and gray boxes indicate unmodified sites.

Considering the extracellular human proteome, only 45 cysteine-containing peptides were identified from vehicle treated samples, 616 and 237 “mucolytic-sensitive” cysteine-containing peptides were detected in P2119- and P2165-treated samples, respectively (Figure 3b). Considering the SARS-CoV-2 proteome, one cysteine-containing peptide was identified in vehicle-treated samples, whereas 15 and 19 “mucolytic-sensitive” cysteine-containing peptides were identified in P2165- and P2119-treated samples, respectively (Figure 3c). In terms of “mucolytic-modified” cysteines, 283 human and 12 viral peptides were identified as covalently modified by P2119. No stable adducts with P2165 were identified in either proteome, as expected. Controls in which viral inactivation was performed addition of urea quench showed a similar trend of peptide identifications (TCEP > P2165 ∼ P2119 > vehicle). Out of 46 cysteine-containing SARS-CoV-2 peptides identified as “mucolytic-sensitive”, the majority were localized to spike protein, *i.e*., 10 out of 15 (67%) peptides after P2165 treatment, 13 out of 19 (68%) peptides after P2119 treatment, and 9 out of 12 (75%) P2119-modified peptides. RBD Cys361, Cys391, Cys432 and Cys525 were susceptible to covalent modification by P2119 (Figure 3d), while disulfides involving Cys432 and Cys525 were sensitive to both P2119 and P2165 (Figure 3e). The peptide containing Cys480 and Cys488 was not identified under any condition implying that accessibility may be a determinant of labeling in the context of the trimeric spike structure. These data indicate that thiol-based mucolytics preferentially target the spike protein relative to other SARS-CoV-2 proteins, and that Cys432 and Cys525 may be distinct from other cysteines in the RBD from the standpoint of reactivity and/or accessibility.

### Mapping mucolytic-sensitive disulfides and reactive cysteines in recombinant SARS-CoV-2 RBD

The goal of subsequent proteomics was to identify “mucolytic-sensitive” disulfides and “hyper-reactive” cysteines in recombinant SARS-CoV-2 RBD (Figure 4a,b and Table 1, Supporting Information) and cross-reference these data to that obtained with SARS-CoV-2. Mass spectrometry (MS) analysis of vehicle-treated RBD identified peptides containing Cys342, Cys361, Cys391, and Cys525, in contrast, while exposure to P2165, P2119 or TCEP control identified peptides containing seven of eight RBD cysteines (Figure 4c) *i.e*., the Cys336 peptide was not found. These mapping data and existing structures therefore suggest that Cys480– Cys488 forms a stable disulfide in recombinant RBD expressed and purified from human HEK293 cells. Notably, this disulfide pair is susceptible to reduction in truncated protein, in contrast to full-length spike harvested from infected cells. The remaining three disulfides Cys336–Cys361, Cys379–Cys432, Cys391–Cys525 exist as an ensemble of free and disulfide-bonded states.

**Figure 4.**
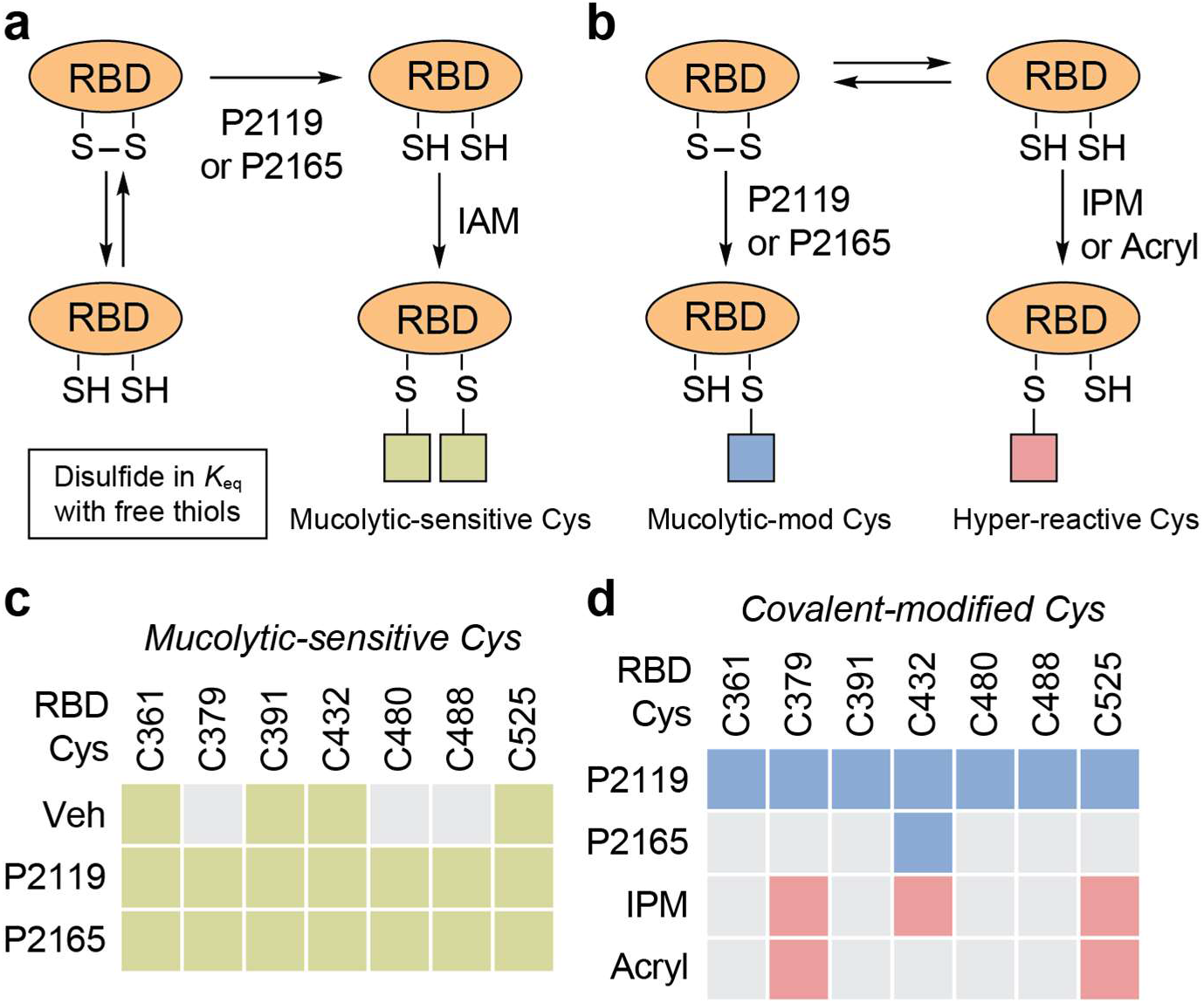
Mucolytic-sensitive, -modified and hyper-reactive cysteines in SARS-CoV-2 RBD. **a**, Workflow for identifying mucolytic-sensitive cysteines in SARS-CoV-2. **b**, Mucolytic-sensitive cysteines in recombinant RBD. Mucolytic-sensitive and unmodified sites are denoted by colored and gray boxes, respectively. **c**, Workflow for identifying mucolytic-modified and hyper-reactive cysteines in SARS-CoV-2. **d**, Mucolytic-modified and hyper-reactive cysteines in recombinant RBD. Mucolytic-modified, hyper-reactive and unmodified sites are denoted by blue, pink, and gray boxes, respectively.

To gain further insight into cysteines reactivity in recombinant RBD, we applied a reactivity-based approach using two different chemical probes for alkylation (Figure 4d). In these studies, RBD was first alkylated with a very low concentration of iodo-*N*-(prop-2-yn-1-yl)acetamide (IPM; 0.1 mM) to detect low p*K*_a_ or “hyper-reactive” cysteines.^33^ Spin gel filtration was used to remove excess IPM and then peptides were incubated with dithiothreitol (DTT). After reduction, nascent thiols were alkylated with a high concentration of IAM (4 mM). MS analysis of peptides derived from these samples revealed that only Cys379, Cys432 and Cys525 were alkylated with IPM. Peptides containing these cysteines also exhibited more intense signals after DTT reduction-IAM alkylation, consistent with powerful thiolate nucleophilicity at these three sites (Figure 4d). By contrast, Cys361, Cys391, Cys480 and Cys488 were only alkylated after reduction with DTT. The reactivities of Cys379, Cys432, and Cys525 were further differentiated using an acrylamide-derivatized electrophile, which is less reactive than IAM and IPM. This compound underwent Michael addition with Cys379 and Cys525, but not Cys432 (Figure 4d) suggesting that inherent nucleophilicity distinguishes these two cysteines in the RBD. Importantly, findings from cysteine reactivity mapping of recombinant SARS-CoV-2 RBD are fully consistent with those obtained from mucolytic-treated samples as well as peptides identified in viral infection studies.

### Computational analysis of disulfide dynamics in the SARS-CoV-2 RBD

Among mobile regions in the RBD is the receptor binding motif (RBM) that includes all RBD amino acids directly participating in the RBD-ACE2 interaction. The RBM is critical, as the plasticity of this motif is related to the ability to recognize and bind ACE2.^34^ Prior studies have characterized residues directly involved in RBD-ACE2 interaction^8,35,36^ and these residues have been parsed into three distinct “contact regions” (CR1–3; Figure 5a).^37^ To dissect the relative contributions of experimentally observed disulfides to dynamics in CR1–3, we conducted MD simulation of the RBD in different redox states. The difference in eigenvector centrality^37^ between native and reduced systems on a per residue basis is depicted in Figure 5b. This analysis facilitates the identification of residues and regions that present significant changes from the mean dynamic behavior in the native state post-disulfide reduction. The resulting data indicate that reduction of the Cys379–Cys432 disulfide bond leads to important dynamic changes in several residues of the interaction region, particularly in CR1. Reduction of the Cys391–Cys525 disulfide was associated with fewer changes overall, but, some residues at CR2 and CR3 were also perturbed. These data were further supported by clustering trajectories using a structure similarity criterion (Figure 5c,d). The RBM region in native RBD was able to access conformations that differed significantly from a reference structure of this domain in complex with ACE2. Reduction of the Cys379–Cys432 disulfide resulted in structure ensembles that differed greatly from the reference structure. Even though reduction of Cys391–Cys525 generated structure ensembles comparable to native RBD in terms of a gross structural descriptor, the observed conformations were quite different. A greater tendency to “shrink” the RBM region approaching CR1 and CR3 was observed, which may have a significant impact on ACE2 recognition; a similar trend was observed for Cys480–Cys488. Although this disulfide pair is located at the RBM, its reduction yielded structure ensembles resembling not only native RBD, but also a subset exploring quite different RBM conformations (Figure S3). In summary, MD simulations suggest that both Cys379–Cys432 and Cys391–Cys525 disulfides, experimentally confirmed by MS as targeted by mucolytic agents P2119 and P2165, play a major role in RBD dynamics. From a mechanistic standpoint, reduction of Cys379–Cys432 and Cys391–Cys525 disulfide pairs appears to impact RBD binding to ACE2, despite being structurally distant from the interaction region.

**Figure 5.**
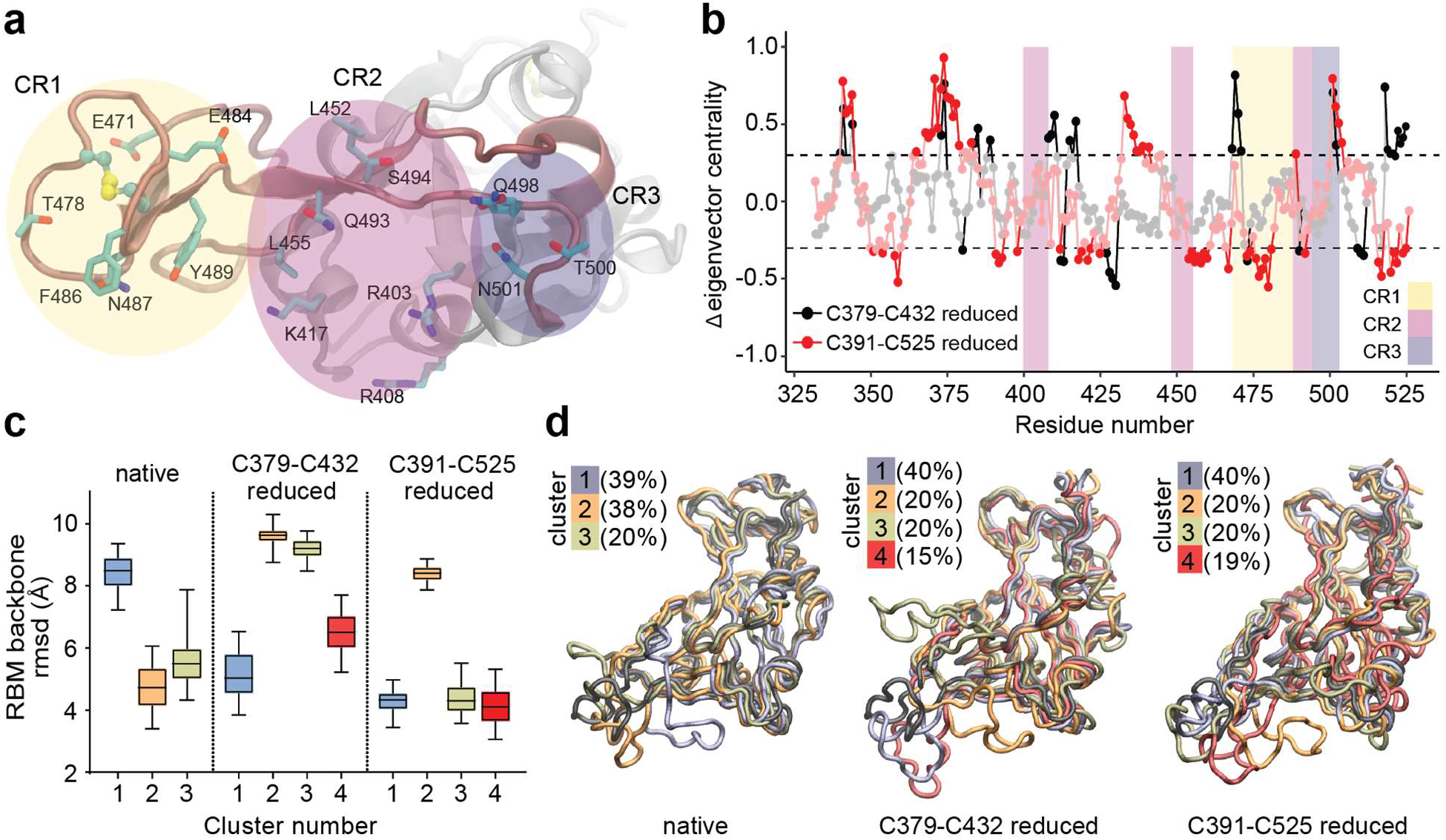
Molecular dynamics simulation of SARS-CoV-2 RBD disulfides. **a**, Detailed representation of RBM residues in RBD that directly interact with ACE2, grouped in three contact regions (CRs). **b**, Eigenvector centrality difference between reduced and native states. CRs regions are colored as in panel a. Residues two sigma units away from normal distribution (dashed line) are shown. **c**, Distribution of the root mean square deviation (rmsd) with respect to the experimental RBM structure for each ensemble of clustered structures. Superposition of the reference RBD structure (black) with a representative RBD structure of each cluster ensemble from different simulations. The percentage of structures belonging to each cluster is indicated.

### Docking thiol-based mucolytics with the RBD of SARS-CoV-2

A hydrophobic binding pocket granting access to the Cys379–Cys432 disulfide was identified in different spike structures at the interface of neighboring RBD subunits. This pocket, shown in Figure 5a, associates with hydrophobic molecules, *e.g*., linoleic acid, even in the closed state of SARS-CoV-2 and related coronavirus spike proteins.^38-40^ The binding of hydrophobic molecules to this pocket has been related to stabilization of spike protein in its closed conformation,^38,41^ and ligands such as poly-unsaturated fatty acids or lipid-soluble vitamins have shown to inhibit SARS-CoV-2 RBD-ACE2 binding *in vitro*.^42^ When analyzing the dynamic behavior of the isolated RBD or its open conformation in the spike protein after extensive MD simulations, this pocket proved to be cryptic, characterized by breathing-like dynamics that allow transient accessibility to the Cys379–Cys432 disulfide (Figure 5 and Figure S4). Available spike structures with hydrophobic ligands bound in this region indicate that larger pocket openings become accessible after binding. Indeed, co-solvent MD simulations from ligand free systems showed that aromatic moieties, such as the benzyl group in P2119 and P2165, can bind *via* induced fit into this intriguing pocket regardless of the glycosylation or conformational state of the RBD.^43^

To explore potential binding modes in greater detail, we evaluated different pocket conformations for docking experiments, including a well-defined pocket due to the presence of a ligand (linoleic acid, PDB 6ZB4) and conformations from our MD simulations (Figure 6a and Table 2, Supporting Information). Using isolated RBD as the receptor template for docking experiments representative of spike open conformations, we found that among the best ranked complex poses were those which allocated P2119 and P2165 to the previously mentioned pocket. In these poses, the benzene moiety of the P2119/P2165 compounds is buried in the protein core, and the thiol points directly towards Cys432 with distances between mucolytic and Cys432 sulfur atoms being less than 5 Å (Figure 6b,c). Several plausible interactions were observed in the docked mucolytic– spike complex. Specifically, the *p*-methoxybenzyl unit forms aromatic interactions with Phe377 and hydrophobic interactions with Leu368/Leu387. The hydrophilic polyol binding is stabilized by polar interactions and hydrogen bonds to RBD backbone carboxyl and amide groups (Figure 6d). Docking poses obtained for both mucolytics were comparable, with only minor differences in the hydroxylated carbon tail conformations. Utilizing RBDs structures derived from the native state MD simulations, in which the pocket has a smaller cavity volume, yielded similar results (Figure 5c,d). Small differences in benzyl orientations were observed, but close distances were consistently maintained between corresponding sulfur atoms (Table 2, Supporting Information). It is worth noting that binding of these mucolytic compounds, as well as other hydrophobic ligands, may rely on the spike open/partially open conformational state, as there are no large channels or cavities connecting this pocket with the protein surface in the closed state of the spike glycoprotein.

**Figure 6.**
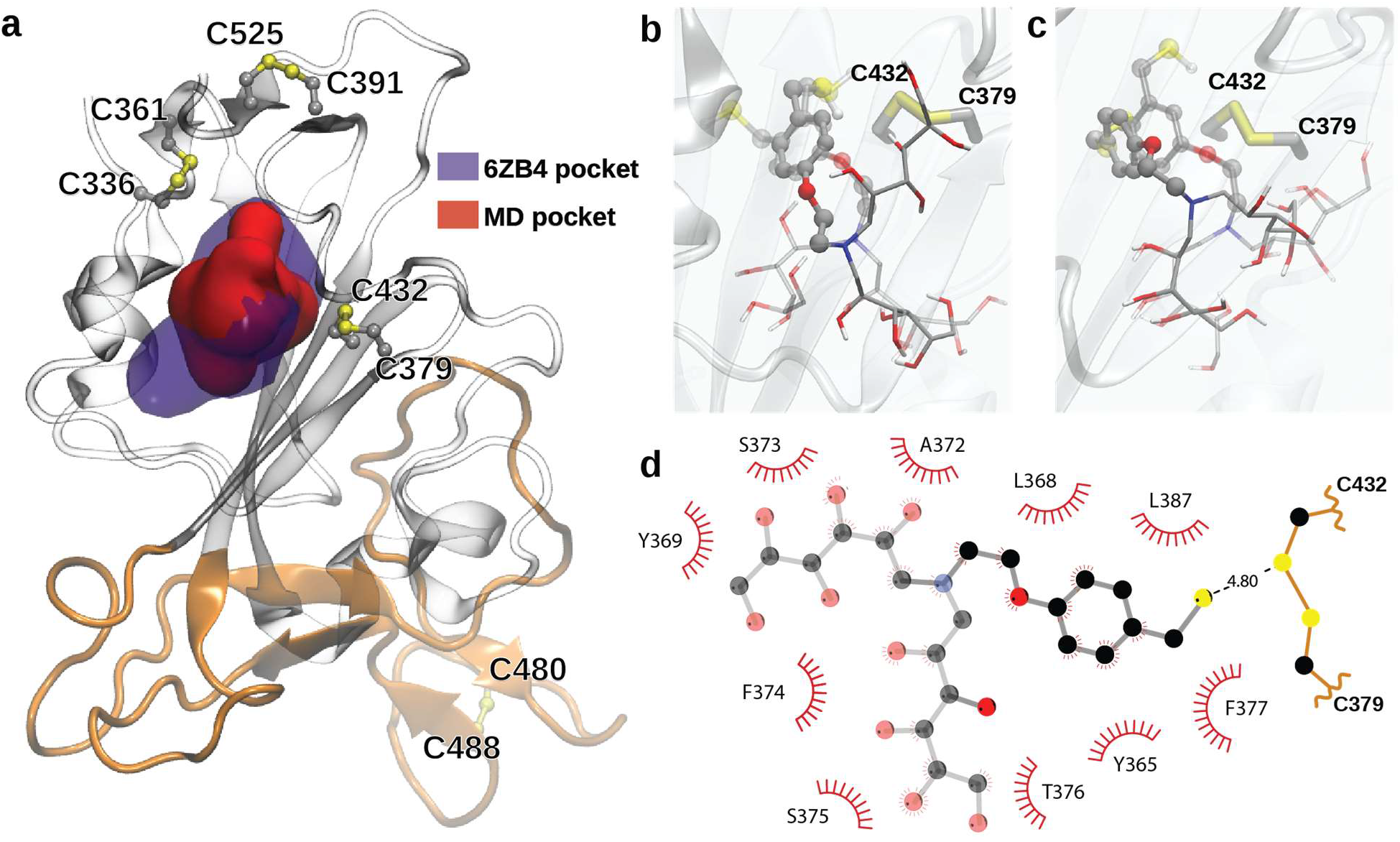
Mucolytics dock in a conserved hydrophobic pocket in the vicinity of C379-C432. **a**, Representation of the hydrophobic pocket in the vicinity of the Cys379-Cys432 disulfide bond. Violet surface corresponds to the pocket identified in the presence of linoleic acid (PDB 6ZB4) and the red surface shows a representative pocket identified during the RBD MD simulations. The RBM is colored orange. **b-c** Superimposed poses of P2119 and P2165 (transparent) docking, selected between the best ranked conformations, using as receptor the RBD of PDB 6ZB4 structure in panel b, or representative conformation from the MD simulation in panel c. **d**, Detailed 2D map highlighting P2119-RBD interactions.

## Discussion

Although several thiol-containing compounds have been shown to inhibit viral receptor binding *in vitro*,^44,45^ they lack potency *e.g*., NAC, glutathione (GSH), or are cytotoxic *e.g*., DTT, TCEP. In this study, we have repurposed mucolytic thiol-based reductants P2119 and P2165 as antiviral agents for human coronaviruses, including SARS-CoV-2. The virucidal effects of these compounds can be attributed to the intrinsic reducing ability of the thiol group, independent of the ability to form stable covalent linkages. Based on their location of action *i.e*., extracellular, the ability of P2119 and P2165 to inhibit RBD-ACE2 binding *in vitro*, and proteomic cysteine site-reactivity mapping in SARS-CoV-2, the simplest explanation for the potent antiviral activity of these mucolytics is that they efficiently reduce key disulfides in the RBD of the SARS-CoV-2 spike protein. The observation that most mapped cysteine modifications occurred on the spike protein supports our proposed mechanism of action for P2119 and P2165. Of particular importance is the reduction of Cys379–Cys432 and Cys391–Cys525, which were identified as “mucolytic-sensitive” in both native spike protein and recombinant RBD. Despite the susceptibility of the Cys480–Cys488 pair to reduction in recombinant RBD, this disulfide was not “in-play” in native virus during infection. In essence, the Cys480–Cys488 disulfide is a conserved, thermodynamically stable disulfide, while Cys379–Cys432 and Cys391–Cys525 disulfides are in dynamic equilibrium with their thiol states and thus, are more sensitive to changes in redox poise.

MD simulations carried out in this study indicate significant conformational changes in the RBM concomitant with reduction of Cys379–Cys432 and Cys391–Cys525 disulfides, even though they are distal to the ACE2 recognition motif. These RBD disulfides appear to be allosteric, a class of disulfides distinct from structural or catalytic disulfides that control protein function by triggering conformational change when they break and/or form.^46^ Experimental data throughout this study also distinguished Cys379–Cys432 and Cys391–Cys525 disulfides including observed mucolytic-dependent changes in cysteine reactivity and mixed disulfide formation with P2119. Furthermore, ligand docking situates P2119/P2165 compounds in similar hydrophobic pockets with access to the Cys379–Cys432 disulfide. Predicted binding poses are supported by the unexpected covalent linkage between Cys432 and P2165 (Figure 4d), likely caused by restricted movement and/or limited solvent access of the resolving thiol in the binding pocket, such that base-promoted ring closure does not occur. Although covalent modification of viral cysteines is not required for the antiviral potency of mucolytic agents, our docking studies outline ostensible interactions between P2119 and P2165, which could be exploited in future work to enhance RBD targeting.

Bjorkman and coworkers recently conducted a structural comparison of SARS-CoV-2 spike neutralizing antibodies (NAbs) and showed NAbs either directly bind to the RBM, or their epitopes include part of the RBM.^47^ For example, a group of NAbs primarily interact with Arg346 and Asn440, which are subject to NAb escape when mutations occur. A recent report by Wilson and coworkers showed that binding and neutralization of the two most common antibody families are abrogated by Lys417Asn and/or Glu484Lys mutations.^48^ Disruption of disulfide bonds adjacent to the aforementioned NAb binding sites *e.g*., Cys336–Cys361 and Cys379–Cys432, could be an alternative mechanism to block ACE2 binding with spike RBD, as evidenced by the potency of P2119 and P2165 against the SARS-CoV-2 Asp614Gly mutant distinguished by an increased level of functional spike, enhanced infectivity, and reinforced ACE2 binding.^49-51^ As SARS-CoV-2 variants, including those with RBM mutations near disulfide bonds like the Glu484Lys and/or Asn501Tyr mutations found in P.1 (gamma), B.1.351 (beta) and B.1.1.7 (alpha) lineages, emerge and pose new threats, targeting spike disulfides offers a variant-independent technique to control the ongoing and future pandemics.

A recent study^52^ suggests that a pro-oxidant environment favoring disulfide formation is required for SARS-CoV-2 to enter host cells *via* the ACE2 receptor and may be the cause behind the age-dependent severity of COVID-19. From this perspective, thiol-based mucolytics P2119 and P2165 may derive antiviral activity from changes in extracellular redox poise in two ways: 1) direct reduction of disulfides on redox-sensitive target proteins required for viral entry *e.g*, RBD Cys379– Cys432 and Cys391–Cys525, and 2) indirectly by restoration of intracellular glutathione levels. Finally, thiol-based mucolytics manage mucus hypersecretion *via* disruption of intermolecular mucin disulfides, with concomitant reduction in airway inflammation and infection.^23^ Accordingly, such agents may be repurposed to alleviate existing SARS-CoV-2 infection in addition to inhibiting viral entry.

### Associated Content Supporting Information

Methods and additional data and figures including virus propagation and inhibition assay, RBD-ACE2 binding assay, LC-MS/MS sample preparation and analysis, docking and molecular dynamics simulations.

## Notes

The authors declare no competing interests. P2119 and P2165 are preclinical pulmonary mucolytics patented (WO2016123335A1) by Parion Sciences, Inc. (Durham, NC, USA).

## Acknowledgements

This work was supported by the US National Institutes of Health (R01 GM102187 and R01 CA174864 to K.S.C, AI089728 and AI110700 to R.S.B.), by grants from Espacio Interdisciplinario (EI_2020) and Comisión Sectorial de Investigación Científica (Grupos_2018), Universidad de la República, to A.Z., S.S. and R.R, and by grants from Fundación Manuel Pérez (Facultad de Medicina, Universidad de la República) and The Richard Lounsbery Foundation to A.Z., S.S., M.R.M. and R.R. Additional support was obtained from Programa de Desarrollo de Ciencias Básicas (PEDECIBA, Uruguay) and Agencia Nacional de Investigación e Innovación (ANII, Uruguay). A.Z., M.R.M. and R.R belong to the SNI program of ANII. This research was also supported by funding from the Chan Zuckerberg Initiative awarded to R.S.B. This project was supported by the North Carolina Policy Collaboratory at the University of North Carolina at Chapel Hill with funding from the North Carolina Coronavirus Relief Fund established and appropriated by the North Carolina General Assembly. The simulations presented in this paper were carried out using ClusterUY (https://cluster.uy).

## References

(1) Cao, B.; Wang, Y.; Wen, D.; Liu, W.; Wang, J.; Fan, G.; Ruan, L.; Song, B.; Cai, Y.; Wei, M.; et al. A Trial of Lopinavir–Ritonavir in Adults Hospitalized with Severe Covid-19. N. Engl. J. Med. 2020, 382 (19), 1787–1799.

(2) Celum, C.; Barnabas, R.; Cohen, M. S.; Collier, A.; El-Sadr, W.; Holmes, K. K.; Johnston, C.; Piot, P. Covid-19, Ebola, and HIV — Leveraging Lessons to Maximize Impact. N. Engl. J. Med. 2020, 383 (19), e106.

(3) Kalil, A. C.; Patterson, T. F.; Mehta, A. K.; Tomashek, K. M.; Wolfe, C. R.; Ghazaryan, V.; Marconi, V. C.; Ruiz-Palacios, G. M.; Hsieh, L.; Kline, S.; et al. Baricitinib plus Remdesivir for Hospitalized Adults with Covid-19. N. Engl. J. Med. 2020, 384 (9), 795–807.

(4) Remdesivir for the Treatment of Covid-19 — Preliminary Report. N. Engl. J. Med. 2020, 383 (10), 992–994.

(5) Beigel, J. H.; Tomashek, K. M.; Dodd, L. E.; Mehta, A. K.; Zingman, B. S.; Kalil, A. C.; Hohmann, E.; Chu, H. Y.; Luetkemeyer, A.; Kline, S.; et al. Remdesivir for the Treatment of Covid-19 — Final Report. N. Engl. J. Med. 2020, 383 (19), 1813–1826.

(6) Hoffmann, M.; Kleine-Weber, H.; Schroeder, S.; Krüger, N.; Herrler, T.; Erichsen, S.; Schiergens, T. S.; Herrler, G.; Wu, N.-H.; Nitsche, A.; Müller, M. A.; Drosten, C.; Pöhlmann, S. SARS-CoV-2 Cell Entry Depends on ACE2 and TMPRSS2 and Is Blocked by a Clinically Proven Protease Inhibitor. Cell 2020, 181 (2), 271-280.e8.

(7) Walls, A. C.; Park, Y.-J.; Tortorici, M. A.; Wall, A.; McGuire, A. T.; Veesler, D. Structure, Function, and Antigenicity of the SARS-CoV-2 Spike Glycoprotein. Cell 2020, 181 (2), 281-292.e6.

(8) Lan, J.; Ge, J.; Yu, J.; Shan, S.; Zhou, H.; Fan, S.; Zhang, Q.; Shi, X.; Wang, Q.; Zhang, L.; Wang, X. Structure of the SARS-CoV-2 Spike Receptor-Binding Domain Bound to the ACE2 Receptor. Nature 2020, 581 (7807), 215–220.

(9) Wang, Q.; Wu, J.; Wang, H.; Gao, Y.; Liu, Q.; Mu, A.; Ji, W.; Yan, L.; Zhu, Y.; Zhu, C.; et al. Structural Basis for RNA Replication by the SARS-CoV-2 Polymerase. Cell 2020, 182 (2), 417-428.e13.

(10) Opstelten, D. J.; de Groote, P.; Horzinek, M. C.; Vennema, H.; Rottier, P. J. Disulfide Bonds in Folding and Transport of Mouse Hepatitis Coronavirus Glycoproteins. J. Virol. 1993, 67 (12), 7394–7401.

(11) Lavillette, D.; Barbouche, R.; Yao, Y.; Boson, B.; Cosset, F.-L.; Jones, I. M.; Fenouillet, E. Significant Redox Insensitivity of the Functions of the SARS-CoV Spike Glycoprotein: Comparison with HIV Envelope. J. Biol. Chem. 2006, 281 (14), 9200– 9204.

(12) Suhail, S.; Zajac, J.; Fossum, C.; Lowater, H.; McCracken, C.; Severson, N.; Laatsch, B.; Narkiewicz-Jodko, A.; Johnson, B.; Liebau, J.; Bhattacharyya, S.; Hati, S. Role of Oxidative Stress on SARS-CoV (SARS) and SARS-CoV-2 (COVID-19) Infection: A Review. Protein J. 2020, 39 (6), 644–656.

(13) Fenouillet, E.; Barbouche, R.; Jones, I. M. Cell Entry by Enveloped Viruses: Redox Considerations for HIV and SARS-Coronavirus. Antioxid. Redox Signal. 2007, 9 (8), 1009–1034.

(14) Lobo-Galo, N.; Terrazas-López, M.; Martínez-Martínez, A.; Díaz-Sánchez, Á. G. FDA-Approved Thiol-Reacting Drugs That Potentially Bind into the SARS-CoV-2 Main Protease, Essential for Viral Replication. J. Biomol. Struct. Dyn. 2020, 1–9.

(15) Debnath, U.; Mitra, A.; Dewaker, V.; Prabhakar, Y. S.; Tadala, R.; Krishnan, K.; Wagh, P.; Velusamy, U.; Subramani, C.; Agarwal, S.; Vrati, S.; Baliyan, A.; Kurpad, V.; Bhattacharyya, P.; Mandal, A. N-Acetyl Cysteine: A Tool to Perturb SARS-CoV-2 Spike Protein Conformation. ChemRxiv 2021, DOI: 10.26434/chemrxiv.12687923.v2

(16) Akhter, J.; Quéromès, G.; Pillai, K.; Kepenekian, V.; Badar, S.; Mekkawy, A. H.; Frobert, E.; Valle, S. J.; Morris, D. L. The Combination of Bromelain and Acetylcysteine (BromAc) Synergistically Inactivates SARS-CoV-2. Viruses 2021, 13 (3), 425.

(17) Jorge-Aarón, R.-M.; Rosa-Ester, M.-P. N-Acetylcysteine as a Potential Treatment for COVID-19. Future Microbiol. 2020, 15 (11), 959–962.

(18) Khanna, K.; Raymond, W.; Charbit, A. R.; Jin, J.; Gitlin, I.; Tang, M.; Sperber, H. S.; Franz, S.; Pillai, S.; Simmons, G.; Fahy, J. V. Binding of SARS-CoV-2 Spike Protein to ACE2 Is Disabled by Thiol-Based Drugs; Evidence from in Vitro SARS-CoV-2 Infection Studies. bioRxiv 2020, DOI: 10.1101/2020.12.08.415505.

(19) Hartl, D.; Starosta, V.; Maier, K.; Beck-Speier, I.; Rebhan, C.; Becker, B. F.; Latzin, P.; Fischer, R.; Ratjen, F.; Huber, R. M.; Rietschel, E.; Krauss-Etschmann, S.; Griese, M. Inhaled Glutathione Decreases PGE2 and Increases Lymphocytes in Cystic Fibrosis Lungs. Free Radic. Biol. Med. 2005, 39 (4), 463–472.

(20) Tirouvanziam, R.; Conrad, C. K.; Bottiglieri, T.; Herzenberg, L. A.; Moss, R. B.; Herzenberg, L. A. High-Dose Oral N-Acetylcysteine, a Glutathione Prodrug, Modulates Inflammation in Cystic Fibrosis. Proc. Natl. Acad. Sci. U. S. A. 2006, 103 (12), 4628 – 4633.

(21) Vasu, V. T.; de Cruz, S. J.; Houghton, J. S.; Hayakawa, K. A.; Morrissey, B. M.; Cross, C. E.; Eiserich, J. P. Evaluation of Thiol-Based Antioxidant Therapeutics in Cystic Fibrosis Sputum: Focus on Myeloperoxidase. Free Radic. Res. 2011, 45 (2), 165–176.

(22) Hancock, L. A.; Hennessy, C. E.; Solomon, G. M.; Dobrinskikh, E.; Estrella, A.; Hara, N.; Hill, D. B.; Kissner, W. J.; Markovetz, M. R.; Grove Villalon, D. E.; et al. Muc5b Overexpression Causes Mucociliary Dysfunction and Enhances Lung Fibrosis in Mice. Nat. Commun. 2018, 9 (1), 5363.

(23) Ehre, C.; Rushton, Z. L.; Wang, B.; Hothem, L. N.; Morrison, C. B.; Fontana, N. C.; Markovetz, M. R.; Delion, M. F.; Kato, T.; Villalon, D.; Thelin, W. R.; Esther, C. R.; Hill, D. B.; Grubb, B. R.; Livraghi-Butrico, A.; Donaldson, S. H.; Boucher, R. C. An Improved Inhaled Mucolytic to Treat Airway Muco-Obstructive Diseases. Am. J. Respir. Crit. Care Med. 2018, 199 (2), 171–180.

(24) Ezeriņa, D.; Takano, Y.; Hanaoka, K.; Urano, Y.; Dick, T. P. N-Acetyl Cysteine Functions as a Fast-Acting Antioxidant by Triggering Intracellular H(2)S and Sulfane Sulfur Production. Cell Chem. Biol. 2018, 25 (4), 447-459.e4.

(25) Li, L.; Rose, P.; Moore, P. K. Hydrogen Sulfide and Cell Signaling. Annu. Rev. Pharmacol. Toxicol. 2011, 51 (1), 169–187.

(26) Hou, Y. J.; Okuda, K.; Edwards, C. E.; Martinez, D. R.; Asakura, T.; Dinnon, K. H.; Kato, T.; Lee, R. E.; Yount, B. L.; Mascenik, T. M.; et al. SARS-CoV-2 Reverse Genetics Reveals a Variable Infection Gradient in the Respiratory Tract. Cell 2020, 182 (2), 429-446.e14.

(27) Zhou, P.; Yang, X.-L.; Wang, X.-G.; Hu, B.; Zhang, L.; Zhang, W.; Si, H.-R.; Zhu, Y.; Li, B.; Huang, C.-L.; et al. A Pneumonia Outbreak Associated with a New Coronavirus of Probable Bat Origin. Nature 2020, 579 (7798), 270–273.

(28) Donaldson, E. F.; Yount, B.; Sims, A. C.; Burkett, S.; Pickles, R. J.; Baric, R. S. Systematic Assembly of a Full-Length Infectious Clone of Human Coronavirus NL63. J. Virol. 2008, 82 (23), 11948 – 11957.

(29) Hofmann, H.; Pyrc, K.; van der Hoek, L.; Geier, M.; Berkhout, B.; Pöhlmann, S. Human Coronavirus NL63 Employs the Severe Acute Respiratory Syndrome Coronavirus Receptor for Cellular Entry. Proc. Natl. Acad. Sci. 2005, 102 (22), 7988– 7993.

(30) Wu, K.; Li, W.; Peng, G.; Li, F. Crystal Structure of NL63 Respiratory Coronavirus Receptor-Binding Domain Complexed with Its Human Receptor. Proc. Natl. Acad. Sci. 2009, 106, 19970–19974.

(31) Goh, J. B.; Ng, S. K. Impact of Host Cell Line Choice on Glycan Profile. Crit. Rev. Biotechnol. 2018, 38 (6), 851–867.

(32) Shaw, D. E. Molecular dynamics simulations related to SARS-Cov-2”. D. E. Shaw Research Technical Data, http://www.deshawresearch.com/resources_sarscov2.html (accessed June 30, 2021).

(33) Fu, L.; Liu, K.; Sun, M.; Tian, C.; Sun, R.; Morales Betanzos, C.; Tallman, K. A.; Porter, N. A.; Yang, Y.; Guo, D.; Liebler, D. C.; Yang, J. Systematic and Quantitative Assessment of Hydrogen Peroxide Reactivity With Cysteines Across Human Proteomes. Mol. Cell. Proteomics 2017, 16 (10), 1815–1828. https://doi.org/10.1074/mcp.RA117.000108.

(34) Raghuvamsi, P. V; Tulsian, N. K.; Samsudin, F.; Qian, X.; Purushotorman, K.; Yue, G.; Kozma, M. M.; Hwa, W. Y.; Lescar, J.; Bond, P. J.; MacAry, P. A.; Anand, G. S. SARS-CoV-2 S Protein:ACE2 Interaction Reveals Novel Allosteric Targets. Elife 2021, 10, e63646.

(35) Shang, J.; Ye, G.; Shi, K.; Wan, Y.; Luo, C.; Aihara, H.; Geng, Q.; Auerbach, A.; Li, F. Structural Basis of Receptor Recognition by SARS-CoV-2. Nature 2020, 581 (7807), 221–224.

(36) Starr, T. N.; Greaney, A. J.; Hilton, S. K.; Ellis, D.; Crawford, K. H. D.; Dingens, A. S.; Navarro, M. J.; Bowen, J. E.; Tortorici, M. A.; Walls, A. C.; King, N. P.; Veesler, D.; Bloom, J. D. Deep Mutational Scanning of SARS-CoV-2 Receptor Binding Domain Reveals Constraints on Folding and ACE2 Binding. Cell 2020, 182 (5), 1295-1310.e20.

(37) Wang, Y.; Liu, M.; Gao, J. Enhanced Receptor Binding of SARS-CoV-2 through Networks of Hydrogen-Bonding and Hydrophobic Interactions. Proc. Natl. Acad. Sci. 2020, 117 (25), 13967 LP – 13974.

(38) Toelzer, C.; Gupta, K.; Yadav, S. K. N.; Borucu, U.; Davidson, A. D.; Kavanagh Williamson, M.; Shoemark, D. K.; Garzoni, F.; Staufer, O.; Milligan, R.; Capin, J.; Mulholland, A. J.; Spatz, J.; Fitzgerald, D.; Berger, I.; Schaffitzel, C. Free Fatty Acid Binding Pocket in the Locked Structure of SARS-CoV-2 Spike Protein. Science 2020, 370 (6517), 725–730.

(39) Zhang, S.; Qiao, S.; Yu, J.; Zeng, J.; Shan, S.; Tian, L.; Lan, J.; Zhang, L.; Wang, X. Bat and Pangolin Coronavirus Spike Glycoprotein Structures Provide Insights into SARS-CoV-2 Evolution. Nat. Commun. 2021, 12 (1), 1607.

(40) Shoemark, D. K.; Colenso, C. K.; Toelzer, C.; Gupta, K.; Sessions, R. B.; Davidson, D.; Berger, I.; Schaffitzel, C.; Spencer, J.; Mulholland, A. J. Molecular Simulations Suggest Vitamins, Retinoids and Steroids as Ligands of the Free Fatty Acid Pocket of the SARS-CoV-2 Spike Protein. Angew. Chemie Int. Ed. 2021, 60 (13), 7098– 7110.

(41) Vivar-Sierra, A.; Araiza-Macías, M. J.; Hernández-Contreras, J. P.; Vergara-Castañeda, A.; Ramírez-Vélez, G.; Pinto-Almazán, R.; Salazar, J. R.; Loza-Mejía, M.A In Silico Study of Polyunsaturated Fatty Acids as Potential SARS-CoV-2 Spike Protein Closed Conformation Stabilizers: Epidemiological and Computational Approaches. Molecules 2021, 26 (3), 711.

(42) Goc, A.; Niedzwiecki, A.; Rath, M. Polyunsaturated ω-3 Fatty Acids Inhibit ACE2-Controlled SARS-CoV-2 Binding and Cellular Entry. Sci. Rep. 2021, 11 (1), 5207.

(43) Zuzic, L.; Samsudin, F.; Shivgan, A. T.; Raghuvamsi, P. V; Marzinek, J. K.; Boags, A.; Pedebos, C.; Tulsian, N. K.; Warwicker, J.; MacAry, P.; Crispin, M.; Khalid, S.; Anand, G. S.; Bond, P. J. Uncovering Cryptic Pockets in the SARS-CoV-2 Spike Glycoprotein. bioRxiv 2021, DOI: 10.1101/2021.05.05.442536.

(44) Grishin, A. M.; Dolgova, N. V; Harms, S.; Pickering, I. J.; George, G. N.; Falzarano, D.; Cygler, M. Spike Protein Disulfide Disruption as a Potential Treatment for SARS-CoV-2. bioRxiv 2021, DOI: 10.1101/2021.01.02.425099.

(45) Manček-Keber, M.; Hafner-Bratkovič, I.; Lainšček, D.; Benčina, M.; Govednik, T.; Orehek, S.; Plaper, T.; Jazbec, V.; Bergant, V.; Grass, V.; Pichlmair, A.; Jerala, R. Disruption of Disulfides within RBD of SARS-CoV-2 Spike Protein Prevents Fusion and Represents a Target for Viral Entry Inhibition by Registered Drugs. FASEB J. 2021, 35 (6), e21651.

(46) Chiu, J.; Hogg, P. J. Allosteric Disulfides: Sophisticated Molecular Structures Enabling Flexible Protein Regulation. J. Biol. Chem. 2019, 294 (8), 2949–2960.

(47) Barnes, C. O.; Jette, C. A.; Abernathy, M. E.; Dam, K.-M. A.; Esswein, S. R.; Gristick, H. B.; Malyutin, A. G.; Sharaf, N. G.; Huey-Tubman, K. E.; Lee, Y. E.; Robbiani, D. F.; Nussenzweig, M. C.; West, A. P.; Bjorkman, P. J. SARS-CoV-2 Neutralizing Antibody Structures Inform Therapeutic Strategies. Nature 2020, 588 (7839), 682– 687.

(48) Yuan, M.; Huang, D.; Lee, C.-C. D.; Wu, N. C.; Jackson, A. M.; Zhu, X.; Liu, H.; Peng, L.; van Gils, M. J.; Sanders, R. W.; Burton, D. R.; Reincke, S. M.; Prüss, H.; Kreye, J.; Nemazee, D.; Ward, A. B.; Wilson, I. A. Structural and Functional Ramifications of Antigenic Drift in Recent SARS-CoV-2 Variants. Science 2021, eabh1139.

(49) Zhang, J.; Cai, Y.; Xiao, T.; Lu, J.; Peng, H.; Sterling, S. M.; Walsh, R. M.; Rits-Volloch, S.; Zhu, H.; Woosley, A. N.; Yang, W.; Sliz, P.; Chen, B. Structural Impact on SARS-CoV-2 Spike Protein by D614G Substitution. Science 2021, 372 (6541) 525–530.

(50) Benton, D. J.; Wrobel, A. G.; Roustan, C.; Borg, A.; Xu, P.; Martin, S. R.; Rosenthal, P. B.; Skehel, J. J.; Gamblin, S. J. The Effect of the D614G Substitution on the Structure of the Spike Glycoprotein of SARS-CoV-2. Proc. Natl. Acad. Sci. 2021, 118 (9), e2022586118.

(51) Plante, J. A.; Liu, Y.; Liu, J.; Xia, H.; Johnson, B. A.; Lokugamage, K. G.; Zhang, X.; Muruato, A. E.; Zou, J.; Fontes-Garfias, C. R.; et al. Spike Mutation D614G Alters SARS-CoV-2 Fitness. Nature 2021, 592 (7852), 116–121.

(52) Giustarini, D.; Santucci, A.; Bartolini, D.; Galli, F.; Rossi, R. The Age-Dependent Decline of the Extracellular Thiol-Disulfide Balance and Its Role in SARS-CoV-2 Infection. Redox Biol. 2021, 41, 101902.

