## Supporting Information for "Thiol-based mucolytics exhibit antiviral activity against SARS-CoV-2 through allosteric disulfide disruption in the spike glycoprotein"

^1^Department of Chemistry, Scripps Research, Jupiter, Florida, USA. ^2^Departamento de Bioquímica, Facultad de Medicina and Centro de Investigaciones Biomédicas (CEINBIO), Universidad de la República, Montevideo, Uruguay.  ^3^Department of Epidemiology, University of North Carolina at Chapel Hill, Chapel Hill, NC, USA. ^4^Institut Pasteur de Montevideo, Montevideo, Uruguay; ^5^Marsico Lung Institute, University of North Carolina at Chapel Hill, Chapel Hill, NC, USA. ^6^State Key Laboratory of Proteomics, Beijing Proteome Research Center, National Center for Protein Sciences • Beijing, Beijing Institute of Lifeomics, Beijing, China. ^7^These authors contributed equally: Yunlong Shi, Ari Zeida, Caitlin E. Edwards. *Corresponding authors:.

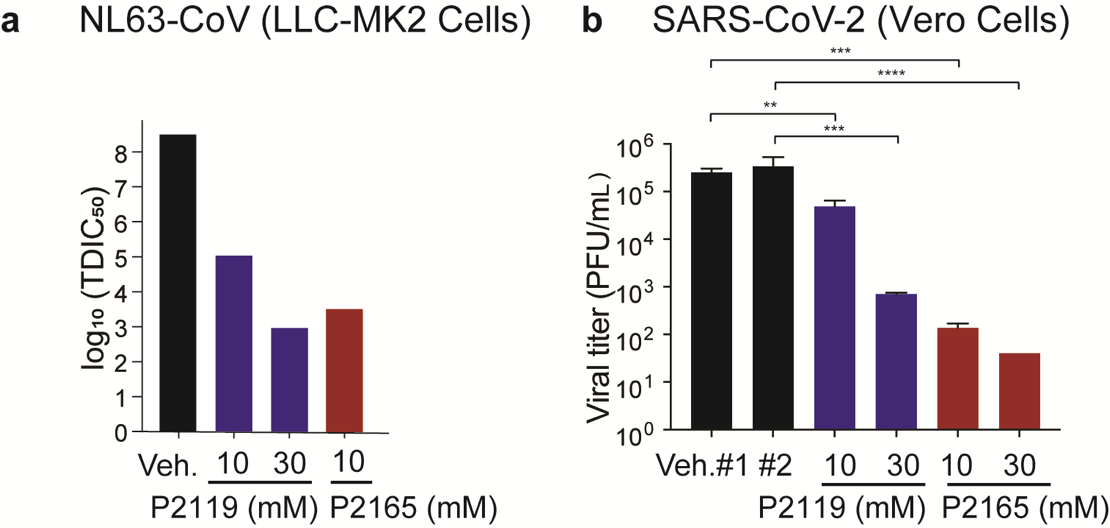

**Figure S1** | **Virucidal activity of thiol-based mucolytics.** **a** A recombinant infectious clone of NL63-CoV was propagated in LLC-MK2 cells. NL63-CoV (100 µl) was incubated with P2165 (10 mM), P2119 (10 mM and 30 mM), or an equal volume of PBS vehicle for 90 min at 37 °C. Virus titers were determined by Median Tissue Culture Infectious Dose (TCID_50_/ml) on Vero cells. **b** Comparison of 72-h titers between vehicle or treated SARS-CoV-2 (D614G) in infected Vero cells at a MOI of 0.1. Triplicated titers of the virus in cultures from the same donor were analyzed by paired t test.

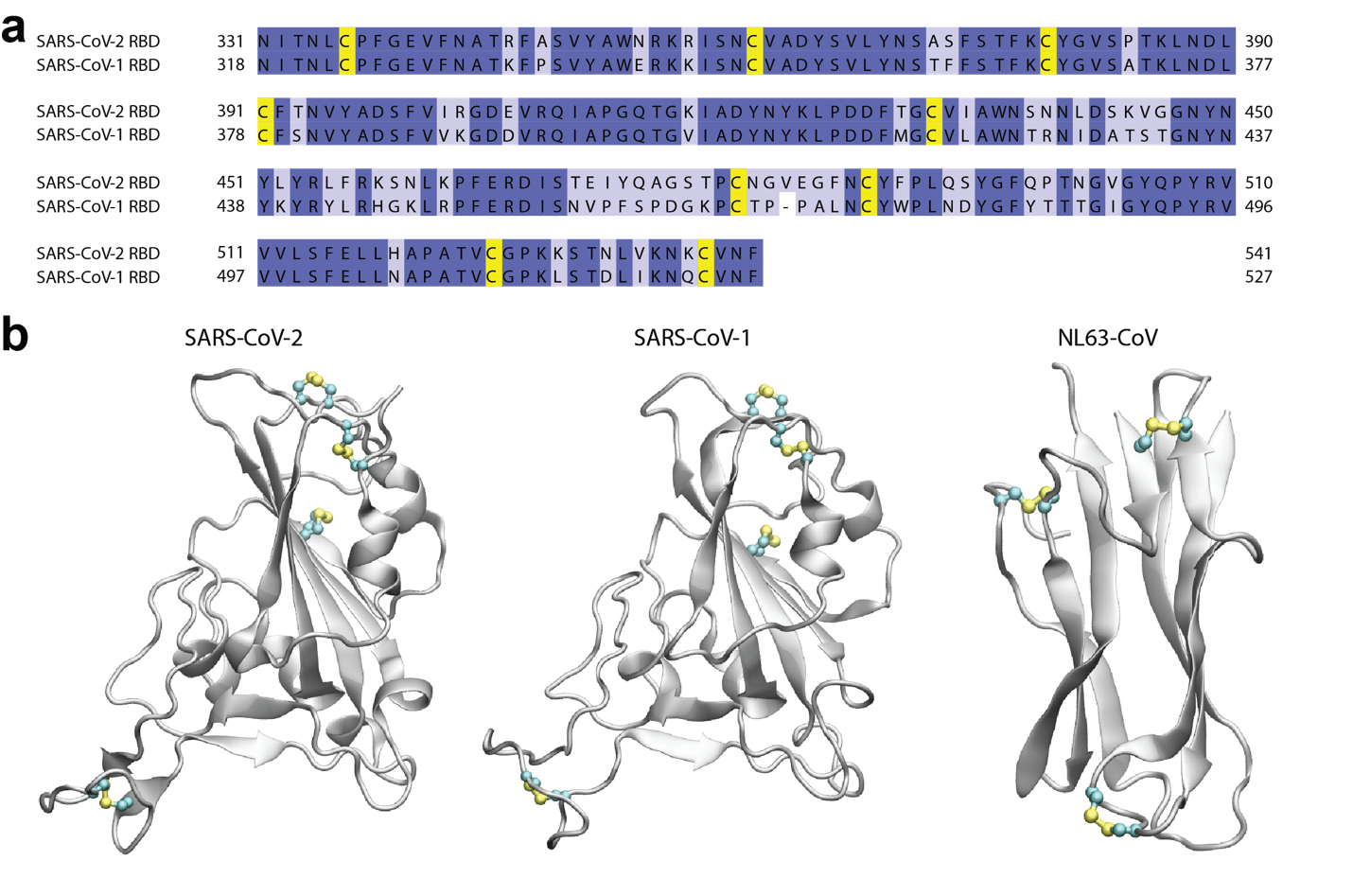

**Figure S2 | Sequence and structure conservation in coronavirus RBD domains. a,** Sequence alignment of SARS-CoV-2 and SARS-CoV-1 RBDs showing fconservation of Cys (yellow). GenBank accession numbers are QHR63250.1 (SARS-CoV-2), AY278488.2 (SARS-CoV-1). **b,** Structural comparison of SARS-CoV-2 (PDB 6LZG), SARS-CoV-1 (PDB 6ACD) and NL63-CoV (PDB 3KBH) RBDs, highlighting native disulfides.

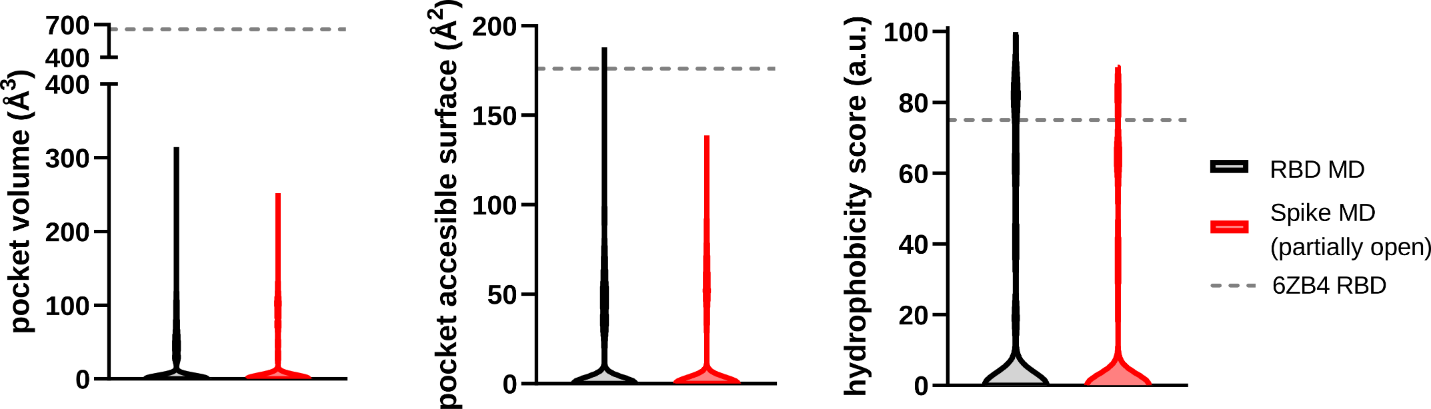

**Figure S3 | Hydrophobic pocket in the vicinity of C379-C432 disulfide bond.** Calculated pocket volume (Å^3^), solvent accessible surface area (Å^2^) and hydrophobicity distributions are shown for the MD simulation of the isolated RBD and also for the simulation of the Spike glycoprotein ectodomain. PDB 6ZB4 structure pocket properties are shown in dashed gray lines as a benchmark.

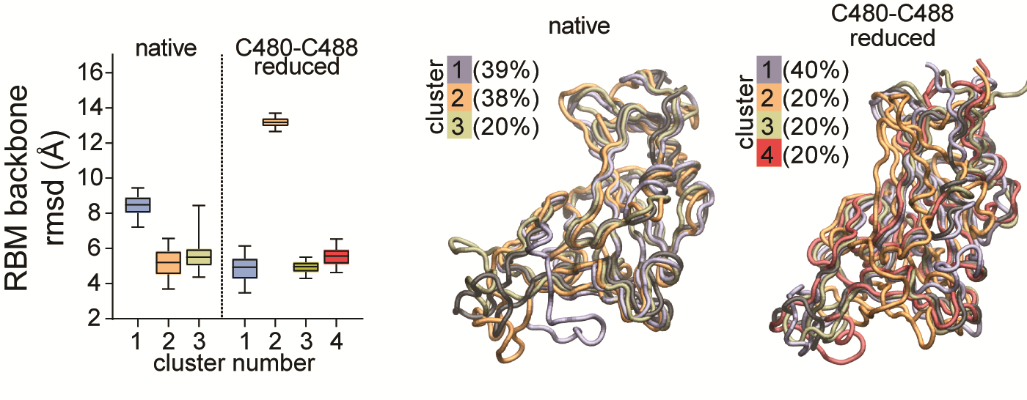
**Figure S4 | Comparison between native and Cys480-Cys488 reduced RBD**. Distribution of the root mean square deviation (rmsd) with respect to the experimental RBM structure for each ensemble of clustered structures. Superposition of the reference RBD structure (black ribbons) with a representative RBD structure of each cluster ensemble from different MDs. The percentage of structures belonging to each cluster is indicated.

**Methods**

**Propagation of SARS-CoV-2 in HNEs and exposure to mucolytic agents*.*** HNE cultures were inoculated at an MOI of 0.1 with 200 µl of SARS-CoV-2 D614G diluted in PBS. Cultures were incubated with virus at 37 °C for 90 min, virus inoculum removed, apical surfaces of cultures washed 3X with 500 µl PBS, and cultures returned to a 37 °C incubator. Apical washes were harvested at 48, 72, and 96 hours post inoculation (hpi). For apical washes, 200 µl PBS was added to the apical surfaces and removed after 10 min at 37 °C and placed in sterile screw-cap tubes. All apical washes from all time-points were pooled into a conical tube and divided into 3 equal aliquots. For the thiol-reducing agent protocols, each aliquot was incubated for 90 min at 37 °C with either P2165 (30 mM), P2119 (30 mM), or equal volume of vehicle (PBS). Following incubation, 16 M urea was added 1:1 with the sample for a final concentration of 8 M urea to inactivate the virus. Samples were incubated for 15 min at room temperature before final storage at 4 °C prior to mass spectroscopy analyses.

**Mucolytic activity against human coronaviruses SARS-CoV, SARS-CoV-2, and NL63-CoV*.*** SARS-CoV-2 (D614G variant) propagated in either HNE cultures or Vero cells was incubated with either P2165 (30 mM), P2119 (30 mM), or vehicle (PBS) for 90 min at 37 °C. Incubated samples were then titered via plaque assay on Vero cells to determine the effects of P-compounds of viral titers.^1^ To compare the antiviral activity of thiol-reducing agents to NAC against both SARS-CoV and SARS-CoV-2 (D614G) both viruses were grown in HNE cultures and virus harvested and pooled as above. Pooled samples of SARS-CoV or SARS-CoV-2 were then incubated with serial dilutions of either P2165 (3, 10, or 30 mM), NAC (10, 30, or 100 mM), or vehicle (PBS) for 90 min at 37 °C and samples titered by plaque assay on Vero cells. Studies on the effects of P-compounds on NL63-CoV infectivity utilized a recombinant infectious clone of NL63-CoV expressing GFP propagated in LLC-MK2 cells.^2^ Briefly, 100 µl of NL63-CoV was incubated with P2165 (10 mM), P2119 (10 mM and 30 mM), or equal volume of PBS vehicle for 90 min at 37 °C and virus titers determined by Median Tissue Culture Infectious Dose (TCID_50_/ml) determined on Vero cells.

**ACE2 receptor binding assay.** The assay was performed with recombinant SARS-CoV-2 spike RBD (SinoBio, Beijing, China) and a commercial ELISA kit (RayBiotech, Peachtree Corners, GA) according to manufacturer’s protocols (n = 4 independent experiments). SARS-CoV-2 spike RBD (0.25 mg/ml in PBS) was treated with PBS, 10 mM TCEP, or 10 mM P2119 for 1 h @ 37 °C. The protein samples were desalted, and diluted in the assay buffer provided by the kit (final concentration 2500 ng/ml). Protein samples (100 µl/well) were serial diluted in assay buffer and incubated in an immobilized ACE2 plate with shaking for 16 h at 4 °C. After washing with kit provided buffer (5X), HRP-conjugated antibody was added and incubated for 1 h at room temperature with shaking. After washing (5X), TMB substrate was added and incubated for 30 min at room temperature, the stop solution added, and the absorbance at 450 nm recorded immediately. EC_50_’s of the binding assay were calculated using the “[Inhibitor] vs response - variable slope (four parameters)” function in Prism (v 7.0.0, GraphPad Software Inc, San Diego, CA).

**Mapping mucolytic-sensitive disulfides in human and SARS-CoV-2 cysteinomes during infection*.*** Proteins from SARS-CoV-2 (D614G) samples were concentrated through precipitation (MeOH-CHCl_3_) or filtering (Amicon Ultra-15, MilliporeSigma, Burlington, MA). After re-dissolving in 50 mM NH_4_HCO_3_ buffer, protein concentration was determined by a BCA assay (Thermo Scientific, Waltham, MA). Next, each sample was diluted to 0.17 mg/ml (50 µl, 8.5 µg proteins, with 0.02% ProteaseMax surfactant), alkylated with IAM (15 mM, 30 min at room temperature in the dark), then digested with sequencing grade trypsin and chymotrypsin (1:40 protease:protein, w/w), acidified with 0.5% TFA, cleaned by C18 Zip-Tip, dried and analyzed by LC-MS/MS.

**Mapping mucolytic-sensitive disulfides and reactive cysteines in recombinant SARS-CoV-2 RBD*.*** SARS-CoV-2 spike RBD expressed in HEK293 cells (SinoBio, Beijing, China, 0.25 mg/ml in PBS) was treated with vehicle (PBS), or TCEP·HCl, P2165 or P2119 (10 mM) for 45 min at 37 °C, followed by 15 min at 55 °C. After cooling to 4 °C, the samples were desalted, buffer exchanged in 50 mM ammonium bicarbonate (pH 8.5) buffer and treated with iodoacetamide (15 mM) at room temperature for 30 min in dark. The samples were digested with chymotrypsin (1:100 ratio, Pierce Biotechnology, Rockford, IL) with ProteaseMax surfactant (0.025%, Promega Corporation, Madison, WI), and cleaned up by C18 ZipTip (2 µg capacity, Millipore, Billerica, MA) according to the manufacturer’s instructions.

**LC-MS/MS analysis.** LC-MS/MS analyses of extracted peptides were carried out using two systems: 1) An Orbitrap Fusion Tribrid mass spectrometer coupled with nEasy-LC1000 nano liquid chromatography (Thermo Fisher Scientific, Waltham, MA), or 2) A QExactive HF-X mass spectrometer equipped with an UltiMate 3000 RSLCnano system (Thermo Fisher Scientific, Waltham, MA). All MS/MS spectra were searched using pFind studio (Ver 3.1.5, Chinese Academy of Sciences).

For an Orbitrap Fusion Tribrid mass spectrometer coupled with nEasy-LC1000 nano liquid chromatography system (Thermo Scientific, San Jose, CA), peptides were eluted from an EASY PepMap^TM^ RSLC C18 column (2 µm, 100Å, 75 µm x 50cm, Thermo Scientific, San Jose, CA) using a gradient of 5-25% solvent B (80/20 acetonitrile/water, 0.1% formic acid) in 45 min, followed by 25-44% solvent B in 15 min, 44-80% solvent B in 0.10 min, a 10 min hold of 80% solvent B, a return to 5% solvent B in 3 min, and finally a 3 min hold of 5% solvent B. The gradient was then extended for the purpose of cleansing the column by increasing solvent B to 98% in 3 min, a 98% solvent B hold for 10 min, a return to 5% solvent B in 3 min, a 5% solvent B fold for 3 min, an increase of solvent B to 98% in 3 min, a 98% solvent B hold for 10min, a return to 5% solvent B in 3 minutes and a 5% solvent B hold for 3 min and finally, another increase to 98% solvent B in 3 minutes and a hold of 98% solvent B for 10min. All flow rates were 250nL/min delivered using a nEasy-LC1000 nano liquid chromatography system (Thermo Scientific, San Jose, CA). Solvent A consisted of water and 0.1% formic acid. Ions were created at 1.9kV using the EASY-Spray^TM^ ion source (Thermo Scientific, San Jose, CA) held at 50^o^C. Data dependent scanning was performed by the Xcalibur v 4.0.27.10 software using a survey scan at 120, 000 resolution in the Orbitrap analyzer scanning mass/charge (m/z) 380-2000 followed by higher-energy collisional dissociation (HCD) tandem mass spectrometry (MS/MS) at a normalized collision energy of 30% of the most intense ions at maximum speed, at an automatic gain control of 1.0E4. Precursor ions were selected by the monoisotopic precursor selection (MIPS) setting to peptide and MS/MS was performed on charged species of 1-8 at a resolution of 30,000. Dynamic exclusion was set to exclude ions after two times within a 30 sec window, for 20 sec.

For the QExactive HF-X system equipped with a UltiMateTM 3000 RSLCnano system (Thermo Scientific, San Jose, CA), peptide sample was loaded onto a fused-silica nano-ESI column (360 µm OD × 150 µm ID) with a needle tip (3~5 µM) packed to a length of 15 cm with a C18 reverse phase resin (AQ 1.9μm, 120Å, ReproSil-Pur). The peptides were separated using an 88min linear gradient from 6% to 95% buffer B (80% MeCN) equilibrated with buffer A (0.1% formic acid) at a flow rate of 600 nL/min across the column. The scan sequence for the Orbitrap began with an MS1 spectrum (Orbitrap analysis, resolution 120,000, scan range of 350-1550 *m/z*, AGC target 3×10^6^, maximum injection time 20ms, dynamic exclusion of 15 seconds). The “Top25” precursors was selected for MS2 analysis, in which precursors were fragmented by HCD prior to Orbitrap analysis ((N)CE 27, AGC target 2×10^4^, maximum injection time 30ms, resolution 15,000, and isolation window: 1.6 Da). All MS/MS spectra were searched using pFind studio (<http://pfind.ict.ac.cn/software/pFind/index.html>). Precursor ion mass and fragmentation tolerance were set as 10 ppm and 20 ppm, respectively. The maximum number of modiﬁcations and missed cleavages allowed per peptide were both set as three. For all analyses, mass shifts of + 15.9949 Da (methionine oxidation) and + 57.0214 Da (iodoacetamide alkylation) and mass shifts caused by the tested chemicals were searched as variable modifications.

**RBD molecular dynamics (MD) simulations.** The initial model used to perform every MD simulation was created using an RBD crystal structure (PDBid 6LZG) to which we have extend both C- and N-terminal regions by using PDBid 6VYB, to a final RBD model from residues 327 to 532 corresponding to the consensus sequence of SARS-CoV-2 lineage B, in which the four disulfide bonds are oxidized (native form). The domain was embedded in an octahedral OPC3 water box^3^ of 97x97x97 Å dimensions. Na^+^ and Cl^-^ ions were added according to the SPLIT method to neutralize and represent a salt concentration of 0.15 M^4^. After minimizing the entire system, heating and equilibration MDs were performed at the NVT and NPT ensembles respectively, during 0.15 ns each. Then, the system was subjected to a three step sampling protocol, consisting first in a 100 ns long plain MD, then 5 independent 200 ns long accelerated molecular dynamics (aMD) were performed, and finally each resulting structure was submitted to a 500 ns long production MD for a total sampling time of 2.5 μs per system. Standard parameters were used for the aMD runs^5^. Production MDs were performed at 300K using the Langevin thermostat^6^ and Berendsen barostat, the SHAKE algorithm was used to keep bonds involving H atoms at their equilibrium length^7^, and periodic boundary and Ewalds sums were used for treating long-range electrostatic interactions. The ff14SB force field^8^ was used for all residues. Every simulation and analysis was performed with the PMEMD and cpptraj modules of the AMBER18 package^9^. All 5 independent MD trajectory replicas were included for analysis.

We also analyzed trajectories of the complete spike glycoprotein ectodomain 10 μs long MDs both in closed and in partially open (1 RBD up) conformations, available to the community by D. E. Shaw Research^10^. Structure visualizations and drawings were performed with VMD 1.9.3^11^ or Protein Imager^12^.

**Docking simulations.** Docking calculations of P2119 and P2165 compounds were performed using standard protocols of AutoDock Vina^13^. Different SARS-CoV-2 RBD conformations were considered as receptor, both coming from our MD simulations, as from PDBid 6ZB4. P2119 and P2165 compounds were prepared with AutoDockTools^14^ by merging nonpolar hydrogen atoms and computing Gasteiger charges. Histidine residues were assumed to be protonated at Nε. The searching grids centers and dimensions are detailed in Table 1, Supporting Information. RBD structures were considered rigid while compounds were allowed to be full flexible. At each run 20 binding modes were generates using an exhaustiveness of 20 and allowing for energy ranges of 100 kcal/mol. Calculations were replicated 500 times for a total of 10,000 poses per ligand. Poses scoring up to 1 kcal/mol from the best found conformation were selected for cluster analysis. Cluster analysis was based on the distance matrix of root mean square deviations (without considering the hydroxylated carbon tails), using a cut-off of 4 Å for grouping poses. On each case, the best pose from the top populated clusters was chosen for analysis.

**References**

(1) Hou, Y. J.; Okuda, K.; Edwards, C. E.; Martinez, D. R.; Asakura, T.; Dinnon, K. H.; Kato, T.; Lee, R. E.; Yount, B. L.; Mascenik, T. M.; et al. SARS-CoV-2 Reverse Genetics Reveals a Variable Infection Gradient in the Respiratory Tract. *Cell* **2020**, *182* (2), 429-446.e14.
(2) Donaldson, E. F.; Yount, B.; Sims, A. C.; Burkett, S.; Pickles, R. J.; Baric, R. S. Systematic Assembly of a Full-Length Infectious Clone of Human Coronavirus NL63. *J. Virol.* **2008**, *82* (23), 11948–11957.
(3) Li, Z.; Song, L. F.; Li, P.; Merz, K. M. Systematic Parametrization of Divalent Metal Ions for the OPC3, OPC, TIP3P-FB, and TIP4P-FB Water Models. *J. Chem. Theory Comput.* **2020**, *16* (7), 4429–4442.
(4) Machado, M. R.; Pantano, S. Split the Charge Difference in Two! A Rule of Thumb for Adding Proper Amounts of Ions in MD Simulations. *J. Chem. Theory Comput.* **2020**, *16* (3), 1367–1372.
(5) Miao, Y.; Feixas, F.; Eun, C.; McCammon, J. A. Accelerated Molecular Dynamics Simulations of Protein Folding. *J. Comput. Chem.* **2015**, *36* (20), 1536–1549.
(6) Pastor, R. W.; Brooks, B. R.; Szabo, A. An Analysis of the Accuracy of Langevin and Molecular Dynamics Algorithms. *Mol. Phys.* **1988**, *65* (6), 1409–1419.
(7) Ryckaert, J.-P.; Ciccotti, G.; Berendsen, H. J. C. Numerical Integration of the Cartesian Equations of Motion of a System with Constraints: Molecular Dynamics of n-Alkanes. *J. Comput. Phys.* **1977**, *23* (3), 327–341.
(8) Maier, J. A.; Martinez, C.; Kasavajhala, K.; Wickstrom, L.; Hauser, K. E.; Simmerling, C. Ff14SB: Improving the Accuracy of Protein Side Chain and Backbone Parameters from Ff99SB. *J. Chem. Theory Comput.* **2015**, *11* (8), 3696–3713.
(9) Case, D. A., *et al.*, Amber 2021, University of California, San Francisco.
(10) Shaw, D. E. Molecular dynamics simulations related to SARS-Cov-2". D. E. Shaw Research Technical Data, <http://www.deshawresearch.com/resources_sarscov2.html> (accessed June 30, 2021).
(11) Humphrey, W.; Dalke, A.; Schulten, K. VMD: Visual Molecular Dynamics. *J. Mol. Graph.* **1996**, *14* (1), 33–38.
(12) Tomasello, G.; Armenia, I.; Molla, G. The Protein Imager: A Full-Featured Online Molecular Viewer Interface with Server-Side HQ-Rendering Capabilities. *Bioinformatics* **2020**, *36* (9), 2909–2911.
(13) Trott, O.; Olson, A. J. AutoDock Vina: Improving the Speed and Accuracy of Docking with a New Scoring Function, Efficient Optimization, and Multithreading. *J. Comput. Chem.* **2010**, *31* (2), 455–461.
(14) Morris, G. M.; Huey, R.; Lindstrom, W.; Sanner, M. F.; Belew, R. K.; Goodsell, D. S.; Olson, A. J. AutoDock4 and AutoDockTools4: Automated Docking with Selective Receptor Flexibility. *J. Comput. Chem.* **2009**, *30* (16), 2785–2791.
